# Hallmarks of uterine receptivity predate placental mammals

**DOI:** 10.1101/2024.11.04.621939

**Authors:** Silvia Basanta, Daniel J. Stadtmauer, Jamie D. Maziarz, Caitlin E. McDonough-Goldstein, Alison G. Cole, Gülay Dagdas, Günter P. Wagner, Mihaela Pavličev

## Abstract

Embryo implantation is essential to pregnancy in eutherian (placental) mammals. The cellular changes that enabled invasive implantation to evolve from superficial embryo apposition, the ancestral condition still retained in marsupials, have remained unclear. We generated and compared single-cell transcriptomes from the peri-implantation and non-pregnant uterus of two rodents with invasive implantation, the mouse and the guinea pig, and the opossum, a marsupial lacking invasive implantation. Cross-species analysis of endometrial cell types and ligand-receptor signaling revealed that the opossum blastocyst-stage uterus, despite lacking fetal-maternal contact, shares key features with the eutherian window of receptivity. These include a window in which the epithelial-stromal *IHH*-*PTCH1* axis is active, but the downstream *BMP2*-mediated signaling that enables decidualization in eutherians is antagonized through a temporary spike in glandular *GREM2*. We propose that a homologous progesterone-responsive endometrial remodeling reaction preceded the evolution of eutherian and marsupial modes of implantation in their respective lineages. Complementary analysis of pre- and post-implantation embryonic transcriptomes from the same species revealed broad conservation of implantation-related functional genes in the blastocyst, as well as a remarkable similarity in ligand production between the mature trophoblast and the luminal epithelium. We suggest that this redundancy between trophectoderm and luminal epithelial signaling was an enabling factor for the evolution of invasive placentation. Together, these findings suggest that eutherian implantation evolved by maternal rewiring of the output of an ancestral window of receptivity-like epithelial-stromal signaling axis toward decidualization, stabilized epithelial loss, and invasion.

**Significance Statement:** Pregnancy in placental mammals begins with implantation and invasion of the embryo, whereas marsupials retain a superficial mode of embryo apposition. How this major evolutionary transition occurred is unknown. Comparing uterine and embryonic single-cell transcriptomes between model rodents and the opossum reveals that the marsupial uterus activates an epithelial-stromal signaling axis canonically associated with receptivity to implantation. In the marsupial uterus, its outputs are opposed by inhibitory signals, whereas in placental mammals, this program promotes uterine decidual transformation and invasion. Early-embryo implantation-related gene expression, in contrast, shows little change across species. These results challenge the view that implantation evolved through the gain of invasive properties by the embryo, and reveal a common developmental foundation from which placentals evolved their novel mode of gestation.

## Introduction

Embryo implantation marks the first sustained cellular interaction between the embryo and the mother during eutherian (placental mammal) pregnancy. In eutherians such as humans and mice, the blastocyst first apposes to the uterine lining, then adheres to the luminal epithelium, and ultimately invades into the endometrial stroma (1). In contrast, marsupials, the most closely related outgroup to eutherian mammals, retain a more superficial mode of pregnancy. Their embryos complete much of development encased in a proteinaceous shell coat, from which it later hatches and forms a superficial placenta that often only lasts 2-3 days and in no known marsupial persists longer than 10 days (2, 3). The evolutionary origin of eutherian implantation was not merely the origin of physical embryo-uterus contact, but the origin of a maternal tissue state permissive to adhesion, epithelial erosion, and stromal invasion.

In eutherians, this permissive state is temporally restricted to the window of receptivity. During this post-ovulatory interval, the uterus undergoes coordinated remodeling, including epithelial expansion through the development of secretory glands, vascular expansion, immune cell recruitment, and pregnancy-associated transformation of endometrial stromal fibroblasts that, in many eutherians including humans and rodents, culminates in decidualization. These changes depend on epithelial-stromal signaling, through which the uterine epithelium integrates cues from ovarian hormones and from the embryo and relays them to the underlying stroma (4, 5). In marsupials, pregnancy also induces endometrial remodeling, classically termed endometrial recognition of pregnancy, including epithelial growth and blebbing and vascular expansion but not luminal epithelial erosion or decidualization (6, 7). Decidualization of the endometrial stroma and erosion of the luminal epithelium are clearly eutherian innovations, and have no counterpart in marsupials (8–11). Nevertheless, whether the upstream cell-signaling cascade that prepares the uterus for implantation was present in some form before invasive implantation originated, and how the evolution of implantation reshaped uterine cell-cell communication, remain important unanswered questions.

The maternal-first model for the evolution of invasive implantation (12) proposes that changes to maternal cell signaling in the eutherian lineage, such as stromal decidualization and luminal epithelial loss, were central drivers. This model predicts that divergent maternal cell signaling patterns and uterine developmental dynamics should be apparent during the transition to early pregnancy in extant marsupials and eutherians. An alternative to this model is the embryo-centric model, which holds that implantation was driven by novel embryonic signals or trophoblast properties that enabled the blastocyst to actively invade the uterine tissue. Distinguishing between these scenarios requires comparing maternal and embryonic cell states across the marsupial-eutherian divide.

Performing this comparison requires establishing the homolog to the eutherian window of receptivity in marsupial gestation. In the gray short-tailed opossum *Monodelphis domestica*, the embryo reaches the blastocyst stage around day 7.5 of a 14.5-day gestation, but remains enclosed within a shell coat and does not directly contact the uterine epithelium (13). Direct fetal-maternal contact begins only after shell coat dissolution on day 12 (14). Because this later post-hatching stage involves direct cell-cell contact between fetal membranes and luminal epithelium (15, 16), luminal epithelial blebbing resembling that observed at eutherian peri-implantation (17), and pro-inflammatory signaling like eutherian post-implantation (18, 19), it has been viewed as the closest marsupial counterpart to eutherian implantation. However, while previous comparison of implantation to post-hatching is based on physical contact and inflammatory responses, it has remained unexplored whether this or an earlier stage in opossum pregnancy exhibits a counterpart to the endometrial remodeling and epithelial-stromal signaling cascade characteristic of the eutherian window of receptivity.

Here, we test these alternatives using comparative single-cell transcriptomics between two rodents and a marsupial. We compare single-cell transcriptomes generated from the non-pregnant and peri-implantation uterus of the mouse *Mus musculus* and the guinea pig *Cavia porcellus*, a non-muroid rodent with interstitial implantation resembling that of humans (20), with paired samples from the opossum non-pregnant and day 7.5 uterus, when the embryo is a shelled blastocyst **(Figure 1a-c)**, and day 13.5 after hatching and during the superficial placentation period sometimes termed implantation (21). We infer ligand-receptor signaling networks, and use differential gene expression analysis to investigate whether and when the marsupial uterus passes through a developmental and cell-signaling state comparable to the eutherian window of receptivity.

**Figure 1.**
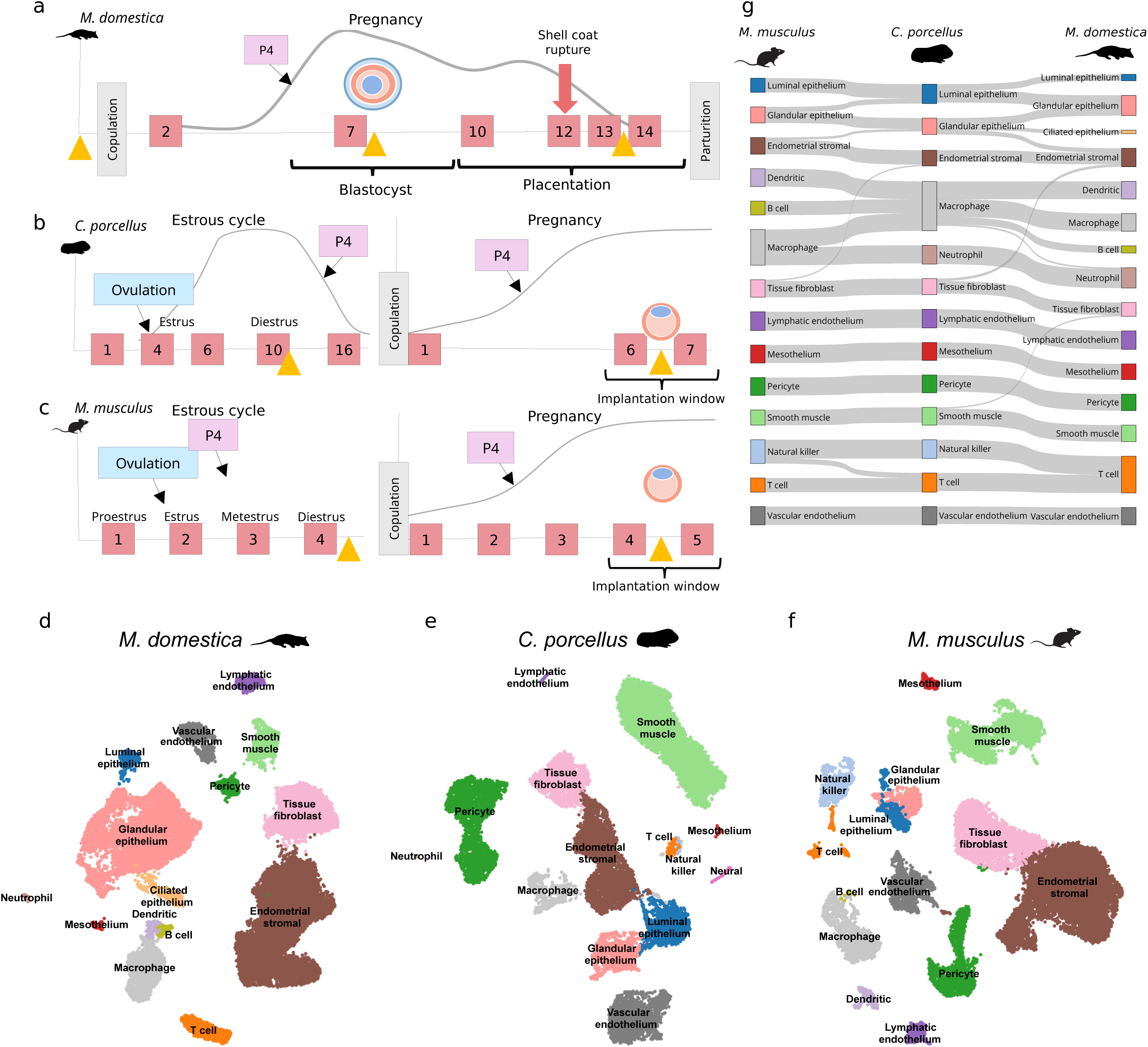
Design of comparable implantation stages across species and resulting cell atlases. **a-c**) Diagrams showing the reproductive cycle timelines of the opossum (**a**) guinea pig (**b**) and mouse (**c**). Yellow arrowheads indicate the stages sampled. Schematic curves of relative maternal serum progesterone are derived from (75–77). P4: progesterone, E2; estradiol. **d-f**) UMAP plots showing cell type annotations for 7.5 dpc opossum (**d**), 6.5 dpc guinea pig (**e**) and 4.5 dpc mouse (**f**). **g**) Sankey plot showing SAMap transcriptomic similarity of cell types across species. Thickness of the bands corresponds to SAMap mapping scores. Mm: *Mus musculus*; Cp: *Cavia porcellus*; Md: *Monodelphis domestica*.

We test for shared gene expression dynamics at the time that the embryo has reached the blastocyst stage, and find that the marsupial uterus presents a transient window of activation similar to the core receptivity-associated epithelial-stromal signaling cascade despite lacking implantation. We investigate the embryo-centric model by comparing time-matched trophectoderm and post-implantation trophoblast transcriptomes from the same species to assess the evolutionary divergence of embryonic gene expression between eutherians and marsupials given the former’s unique invasive capacity and luminal epithelial loss. These analyses allow us to pick apart three separate evolutionary puzzles that are easily conflated due to their origin along the same branch of the mammalian phylogeny: how the window of uterine receptivity evolved, how eutherians acquired invasive implantation, and whether the decisive changes occurred in the embryo or in maternal tissue. Based on these results, we consider an evolutionary model in which conserved progesterone-responsive signaling axis governing uterine remodeling preceded the divergence of marsupials and placental mammals, and argue that eutherian implantation evolved by modifying the output from this ancestral maternal program.

## Results

### Cell type composition across species

Single-cell RNA sequencing of non-pregnant and peri-implantation or equivalent uteri from mouse, guinea pig, and opossum yielded 77,829 high-quality cells after filtering low-count cells and predicted doublets **(Figure S1)**. Leiden clustering identified 14-15 major cell types per species, which were annotated by established markers from single-gene expression studies and comparison to previous mouse and opossum atlases **(Figure 1d-f) (Table S1**) (21, 22). Projection into a shared cross-species embedding using SAMap (23) showed strong correspondence among annotated cell types **(Figure S2; Table S2)**, with most pairwise comparisons yielding unambiguous homologous assignments (**Figure 1g**).

Major epithelial and stromal cell types present at that stage of pregnancy were conserved across species. Epithelial cells expressed canonical markers including *EPCAM*, *MSX2*, *KLF5*, and *WFDC2*, and separated into transcriptomically distinct luminal epithelial (LE) and glandular epithelial (GE) populations in all three species (**Figure 1d-f; Figure S3**). Glandular epithelial cells in all species showed expression of *FOXA2* and *ELAPOR1*, whereas luminal epithelial cells were enriched for *PAX8*, *WFDC2*, and the stem cell marker *LGR5* (24). In the opossum, we also recovered ciliated epithelial (CE) cell population enriched in *FOXJ1* and *ADGB* (25) that mapped most closely to glandular epithelium of the other species (**Figure 1g**) and was not detected in either rodent, consistent with histological evidence that ciliated cells are restricted to the oviduct in rodents (26, 27). Luminal and glandular epithelial cell types mapped specifically across species (**Figure 1g**), supporting consistency of these cell type identities across the marsupial-eutherian divide.

Fibroblasts shared expression of *LUM* and consistently clustered into two cell populations, endometrial stromal fibroblasts (ESF) and myometrial tissue fibroblasts (TF) **(Figure S3)**. ESF were enriched for *HOXA11*, *NR2F2*, *PGR,* and *SMOC2*, whereas TF expressed markers including *CLEC3B*, *DPT*, *ECM1,* and *FBLN1* **(Figure S3)**. Previous studies have shown that these populations are associated with different tissue microenvironments: ESF primarily represent subepithelial and peri-glandular fibroblasts, whereas TF lie closer to the myometrium in both eutherians and the opossum (21, 28). Thus, principal epithelial and stromal cell types were identifiable and comparable across the three species.

### Marsupial uterine remodeling expands glands, whereas rodent receptivity is stromal

To characterize change in the relative cell type composition of the uterus during the transition to the peri-implantation period, we compared the relative proportions of broad cell type classes (epithelial, stromal, vascular, etc.) across the stages sampled. In all three species, stromal cells, primarily fibroblasts, comprised greater than 50% of the captured cells in the non-pregnant state **(Figure 2a)**. Epithelial cell abundance was elevated in early pregnancy, and was most pronounced in the opossum, where epithelial cells after day 7.5 exceeded 50% of all cells captured (**Figure 2a**). Glandular epithelial cells were largely responsible for this change (**Figure 2b**), consistent with quantitative histology documenting a marked increase in gland area coinciding with the post-ovulatory hormone surge (29). Conversely, fibroblast cell types showed a greatly reduced abundance in the pregnant opossum uterus not observed in the two eutherian species **(Figure 2c**).

**Figure 2.**
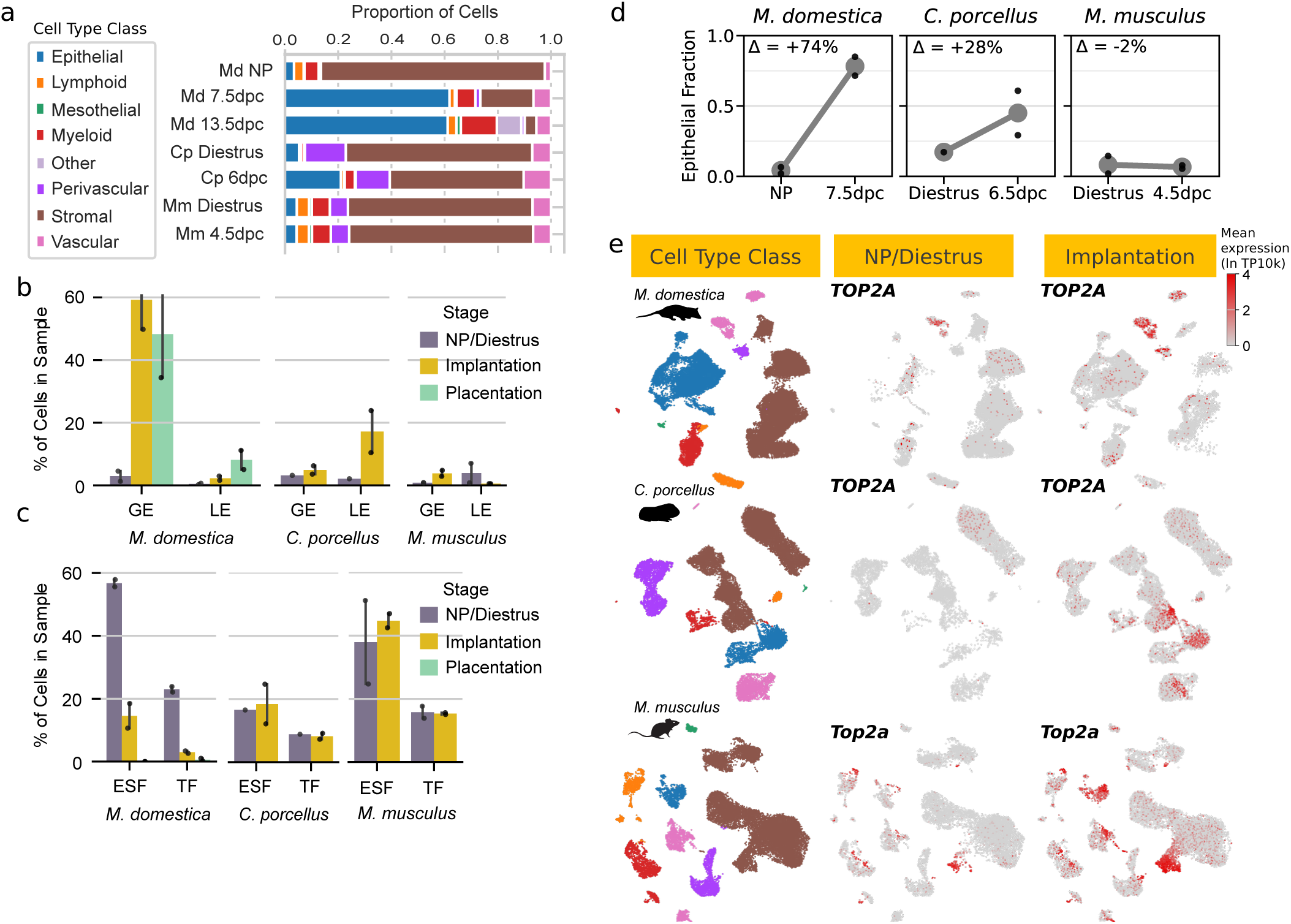
Changes in cell abundance and relative proliferation at peri-implantation. **a)** Relative abundance of cell type classes across species and stages. Md: *Monodelphis domestica*; Cp: *Cavia porcellus*; Mm: *Mus musculus*; NP: non-pregnant. **b**) Barplot abundance of glandular (GE) and luminal (LE) epithelial cells divided by stage and species. Error bars represent the range. NP: non-pregnant. **c**) Barplot showing stromal cell type abundance across species. ESF: endometrial stromal fibroblast; TF: tissue fibroblast. d) Epithelial fraction, defined as the number of epithelial cells captured divided by the sum of epithelial and fibroblast cells captured, for all species. e) UMAP plots showing the distribution of cell type classes (left, colored as in (**a**)) and expression of proliferation marker TOP2A at non-pregnant (center) and peri-implantation (right) timepoints in opossum (top), guinea pig (center), and mouse (bottom).

Cross-species comparison of the epithelial-stromal fraction - defined as the number of epithelial cells captured divided by the total number of epithelial cells and fibroblasts - revealed different temporal trajectories across species. In the opossum, the epithelial fraction greatly expanded at the blastocyst-stage uterus compared to non-pregnant uterus (+74%), whereas changes in guinea pig (+28%) were more moderate and absent in the mouse (-2%) **(Figure 2d)**. Endometrial stromal fibroblasts showed increased expression of the cell cycle marker *TOP2A* **(Figure 2e)** in both the mouse (from 54 to 261 TPM, FDR = 9.1 x 10^-37^) and guinea pig (from 0 to 116 TPM, FDR = 2.3 x 10^-11^), indicating that the stroma proliferates in early rodent pregnancy.

Pronounced epithelial expansion in the opossum raised the question of which developmental programs underlie this remodeling. In eutherians, uterine gland remodeling involves separable morphogenetic processes of epithelial growth and crypt morphogenesis. Estradiol-responsive growth factors, especially IGF1, are among the best-established mediators of uterine epithelial proliferation (30, 31), whereas Wnt signals, including WNT5A and WNT7A, have been implicated in budding and invagination of new crypts from the lumen (24, 32, 33). We therefore compared these two pathways across species. Glandular epithelial cell *IGF1* expression significantly increased at the blastocyst stage in all three species (**Figure 3a**), and the greatest increase by far was in the opossum, where glandular epithelial *IGF1* rose from 7 to 2076 TPM (FDR = 2.9 x 10^-26^). To test whether this corresponded with a transcriptome-wide signature of estradiol response, we conducted gene set enrichment analysis on the full set of genes upregulated in the opossum glandular epithelial cells between non-pregnant and 7.5 dpc states using the Enrichr “Gene_Pertubations_from_GEO_up” database, which catalogs experimental hormone, cytokine, and growth factor perturbations on cell lines and animal models This analysis revealed significant enrichment in estradiol-related pathways, including “estradiol mouse BAL cells” and “17beta-estradiol mouse uterus”, as well as growth pathways “growth hormone mouse adipocytes” and “insulin-like growth factor I, human MCF-7 cells” **(Figure 3b)**. These results are consistent with a mechanism where estrogen-induced activation of *IGF1* drives glandular expansion and shifts the epithelial-stromal ratio towards an epithelial-dominated state (34). Signaling related to gland expansion peaks earlier in the cycle around the shelled blastocyst stage, after which the expanded glands persist and respond to fetal cytokine signaling (35), but no longer express high levels of *IGF1* (**Table S3**).

**Figure 3.**
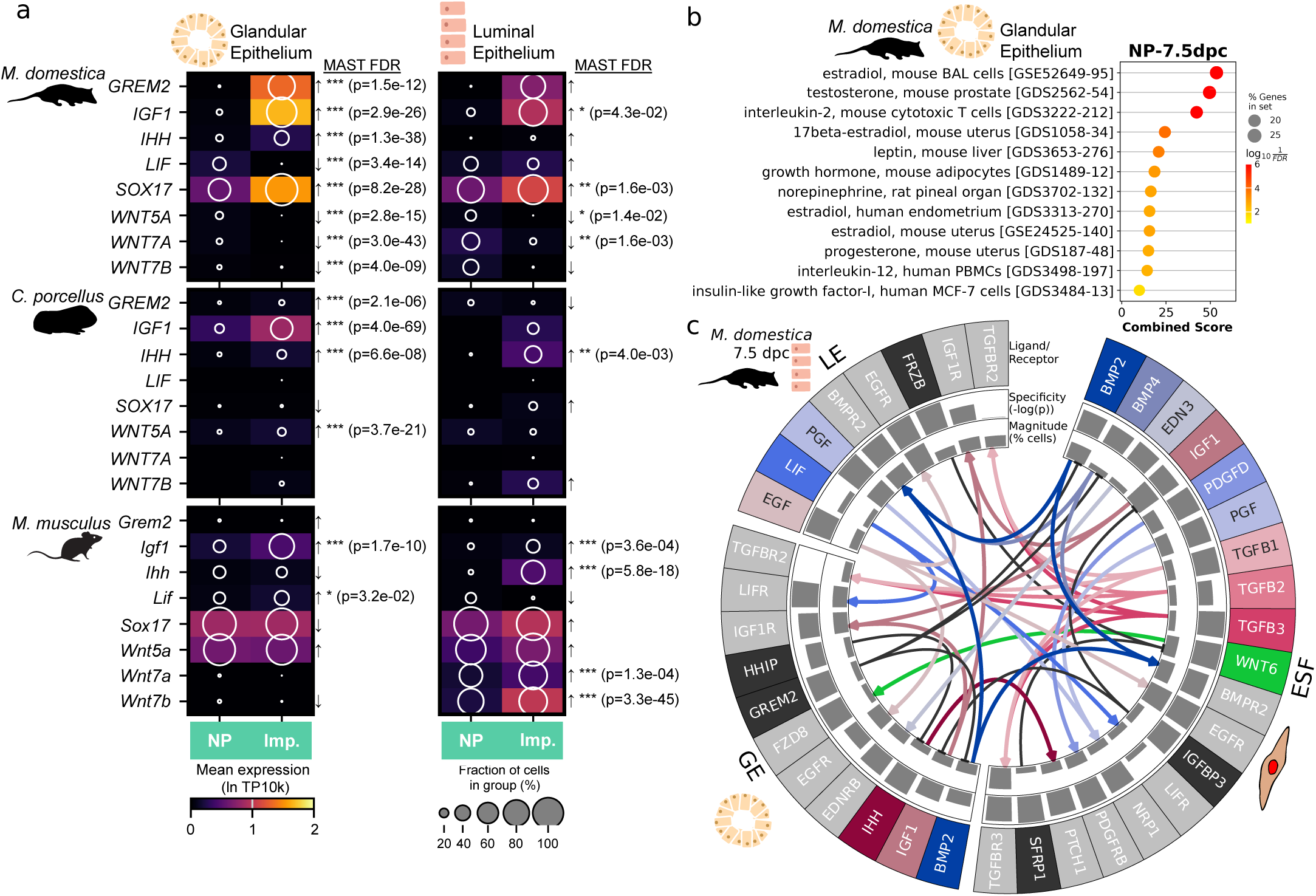
Gene expression changes and epithelial-stromal ligand-receptor interactions at peri-implantation. **a**) Dot plots of implantation-relevant developmental markers showing expression levels and fractions of cells in each cluster. Up and down arrows represent positive and negative fold-change differences between the two stages. *: FDR < 0.05; **: FDR < 0.01; ***: FDR < 0.001. **b**) Statistical scoring from GSEApy gene set enrichment analysis on differentially expressed genes in the opossum glandular epithelial cells between non-pregnant and 7.5 dpc using the EnrichR “Gene Perturbations from GEO up” database as the test gene set. **c**) Inferred ligand-receptor signaling network between select epithelial and stromal cell types in the opossum day 7.5 uterus. Arrows show directions of inferred ligand-receptor signaling. Outer barplots show LIANA+ rank-aggregate specificity score, while inner barplots show the percent of cells in the cluster expressing the ligand or receptor. Ligands are shown in color, receptors in grey, and inhibitors in black. ESF: endometrial stromal fibroblast; GE: glandular epithelial cell; LE: luminal epithelial cell.

We next asked whether the blastocyst-stage endometrium showed activation epithelial Wnt signaling. In the mouse, several Wnt ligands were significantly upregulated at peri-implantation, including luminal epithelial *Wnt7a* (from 23 to 61 TPM, FDR = 1.3 x 10^-4^), *Wnt7b* (from 32 to 353 TPM, FDR = 3.3 x 10^-45^), and *Wnt11* (from 7 to 131 TPM, FDR = 6.3 x 10^-15^), and a positive trend in *Wnt5a* (from 55 to 109 TPM, FDR = 0.23). Guinea pig luminal epithelial cells showed a significant increase in *WNT5A* (from 16 to 31 TPM, FDR = 3.7 x 10^-21^) and positive trends in expression of *WNT7B* (from 65 to 86 TPM, FDR = 0.20) and *WNT11* (from 4 to 38 TPM, FDR = 0.68). In contrast, Wnt ligands showed the opposite trend in the opossum, and decreased in expression at day 7.5 (*WNT7A* from 44 to 18 TPM, p = 1.6 x 10^-3^; *WNT7B*: from 19 to 2 TPM, p = 0.14; *WNT5A*: from 9 to 0 TPM, p = 6.96 x 10^-13^; *WNT11*: from 59 to 5 TPM, p = 2.1 x 10^-4^), and remained low at day 13.5 (**Table S3**). Overall, whereas rodent peri-implantation showed concurrent *IGF1* and epithelial Wnt signaling, the opossum blastocyst-stage uterus showed strong estradiol/*IGF1*-associated glandular expansion despite reduced Wnt ligand expression. These data suggest that marsupial epithelial remodeling at this stage is dominated by glandular growth and expansion but not activation of the Wnt pathway observed in rodents.

### The epithelial-stromal IHH signaling axis is conserved between the eutherian window of implantation and marsupial pre-attachment

Most knowledge about mechanisms of implantation derives from mouse models. In the mouse uterus, cellular changes during the window of receptivity arise from the exchange of morphogens between uterine epithelial and stromal cells under the temporal coordination of sex hormones (36). To further investigate the evolution of these epithelial-stromal signaling patterns, we used LIANA+ (37) to infer ligand-receptor signaling potential between epithelial and stromal cells from cell type transcriptomes, and used differential expression testing between non-pregnant and blastocyst stages to detect temporal trends **(Figure 3; Table S3; Table S4)**.

Epithelial-stromal signaling via the Indian hedgehog (IHH)-PTCH1 axis was inferred in the peri-implantation stage of the two eutherian species and during the blastocyst stage of the opossum (**Figure 3c-e**). IHH signaling from the epithelium to the stroma is essential for uterine receptivity (38). In the mouse, luminal epithelial cell expression of *IHH* increased from 6 to 101 TPM (FDR = 5.8 x 10^-18^), and in the guinea pig increased from 8 to 123 TPM (FDR = 4.0 x 10^-3^) (**Figure 2h**). In the opossum, *IHH* showed significantly increased expression in glandular epithelial cells at the blastocyst stage (from 9 to 45 TPM, FDR = 1.3 x 10^-38^), whereas the change in luminal epithelial expression was less pronounced (0.5 to 3 TPM, FDR = 0.14). In rodents, expression of *IHH* has been shown to be regulated by progesterone, which binds epithelial PGR and induces expression of the transcription factor *SOX17*, which in turn drives *IHH* expression (39–41). Indeed, *SOX17* expression was significantly increased in the opossum luminal and glandular epithelial cells concomitant with *IHH* expression (**Figure 3a**), consistent with its regulation through the same mechanism. Furthermore, we compared stromal receptor expression, as cell signaling strength is a product of both ligand and receptor expression levels. In both guinea pig and opossum, expression of the complementary receptor *PTCH1* was greatest in ESF, and increased at the blastocyst stage (from 83 to 330 TPM in guinea pig, FDR = 1.3 x 10^-57^; from 121 to 202 TPM in opossum, FDR = 1.2 x 10^-37^). In contrast, at the 13.5 dpc placentation stage of the opossum, expression of *IHH* in epithelial cells and *PTCH1* in ESF cells were both absent **(Table S3)**, suggesting only a transient activation at the shelled blastocyst stage. These findings suggest that the progesterone-driven epithelial-stromal IHH-PTCH1 axis is conserved during a limited window of therian mammal early pregnancy that corresponds to the eutherian window of implantation.

Lastly, because LIF is a central regulator of implantation in the mouse (42), we asked whether LIF signaling showed a similar conserved pattern. In the mouse, peri-implantation estrogen induces glandular expression of *Lif*, which activates LIFR-IL6ST/STAT3 in the luminal epithelium to promote receptivity-associated epithelial differentiation and downstream implantation pathways, including IHH (5, 43, 44). Indeed, *Lif* was expressed and increased at peri-implantation in glandular epithelial cells in the mouse (from 13 to 33 TPM, FDR = 3.2 x 10^-2^) (**Figure 3a**). In the opossum, however, *LIF* expression decreased in the glandular epithelium (from 49 to 0.5 TPM, FDR = 3.4 × 10^-14^), along with its receptor *LIFR* in the luminal epithelium (from 54 to 8 TPM, FDR = 0.23). In the guinea pig, *LIF* was not detected in glandular or luminal cells at the sampled peri-implantation stage. Thus, unlike the IHH-PTCH1 axis, *LIF* did not show a consistent activation pattern across the marsupial-eutherian divide or even between the mouse and guinea pig, and we therefore interpret LIF as a lineage-variable component of receptivity outside of the core ancestral epithelial-stromal program.

### Stromal cells show opposing dynamics in eutherians and marsupials, possibly linked to marsupial-specific BMP antagonism

An essential downstream effect of peri-implantation IHH in eutherians is decidualization of endometrial fibroblasts. In eutherians, the epithelial IHH-stromal PTCH1 signaling axis is necessary for decidualization through a BMP2-dependent pathway in stromal cells shown to activate the transcription factor NR2F2 (COUP-TFII) (45–48). *NR2F2* was highly expressed (>300 TPM) in ESF of all species, but did not show a temporal trend (**Table S2**). Investigation of the BMP2 signaling pathway, however, revealed a temporal and species-specific difference. The blastocyst-stage ligand-receptor signaling network of the opossum suggested inhibition from glandular epithelial cell *GREM2*, a secreted antagonist of BMP2 and BMP4 signaling (**Figure 3a**; **Figure 3c**) (49). *GREM2* is greatly upregulated at the blastocyst stage in glandular epithelial cells (from 2 to 2134 TPM, FDR = 1.5 x 10^-12^). Cell-cell signaling analysis inferred that although stromal fibroblast *BMP2* and *BMP4* expression is present at this stage of the opossum, glandular epithelial cell *GREM2* may inhibit it, in parallel with *HHIP* production that antagonizes upstream IHH as well **(Figure 3a)**. *GREM2* returned to low expression (<10 TPM) in 13.5 dpc uterine glands, suggesting a temporary spike around the blastocyst stage. As BMP2 is a key output from the IHH-PTCH1 axis responsible for stromal decidualization in rodents, the presence of BMP2 antagonism in the opossum suggests that while progesterone-IHH-PTCH1 axis itself is conserved, its downstream sequelae may be cut off at the point of stromal BMP2.

In many eutherians including rodents, the decidualization reaction only completes after embryo contact; in our peri-implantation time points, decidualization had not yet occurred. We nevertheless investigated whether common early steps of ESF transformation characterize the rodent window of receptivity before embryo contact is made, and whether they have a counterpart in the non-deciduate opossum. Direct comparison of transcriptomes from a single time-point is subject to bias from lineage-specific transcriptomic properties. Therefore, we instead calculated the overlap of differentially expressed gene sets between the non-pregnant and blastocyst stage ESF of all species using hypergeometric overlap analysis (50), a design that targeted gene expression programs changing in the same (concordant) or opposite (discordant) direction at the window of receptivity. ESF of the mouse and guinea pig displayed a high concordance of gene expression changes with one another, including both up- and down-regulated genes (**Figure 4a**). In contrast, opossum ESF showed a combination of gene expression changes in the same direction as the two eutherians, and discordant changes most concentrated in the upper-left quadrant (**Figure 4b-c**), i.e. upregulated in rodent early decidualization but down-regulated in opossum ESF. Finally, we compared the ESF gene expression changes between non-pregnant and the 13.5 dpc placentation stage of the opossum, when few stromal cells remain, likely due to apoptosis (51): in this case, the direction of change was reversed relative to the rodent transition to stromal receptivity **(Figure 4d-e)**. Overall, these comparisons, while consistent with the opossum shelled blastocyst stage being the homolog to the eutherian window of receptivity, reveal that opossum ESF differ from eutherian ESF in their reaction to the endocrine and paracrine signaling environment of this stage.

**Figure 4.**
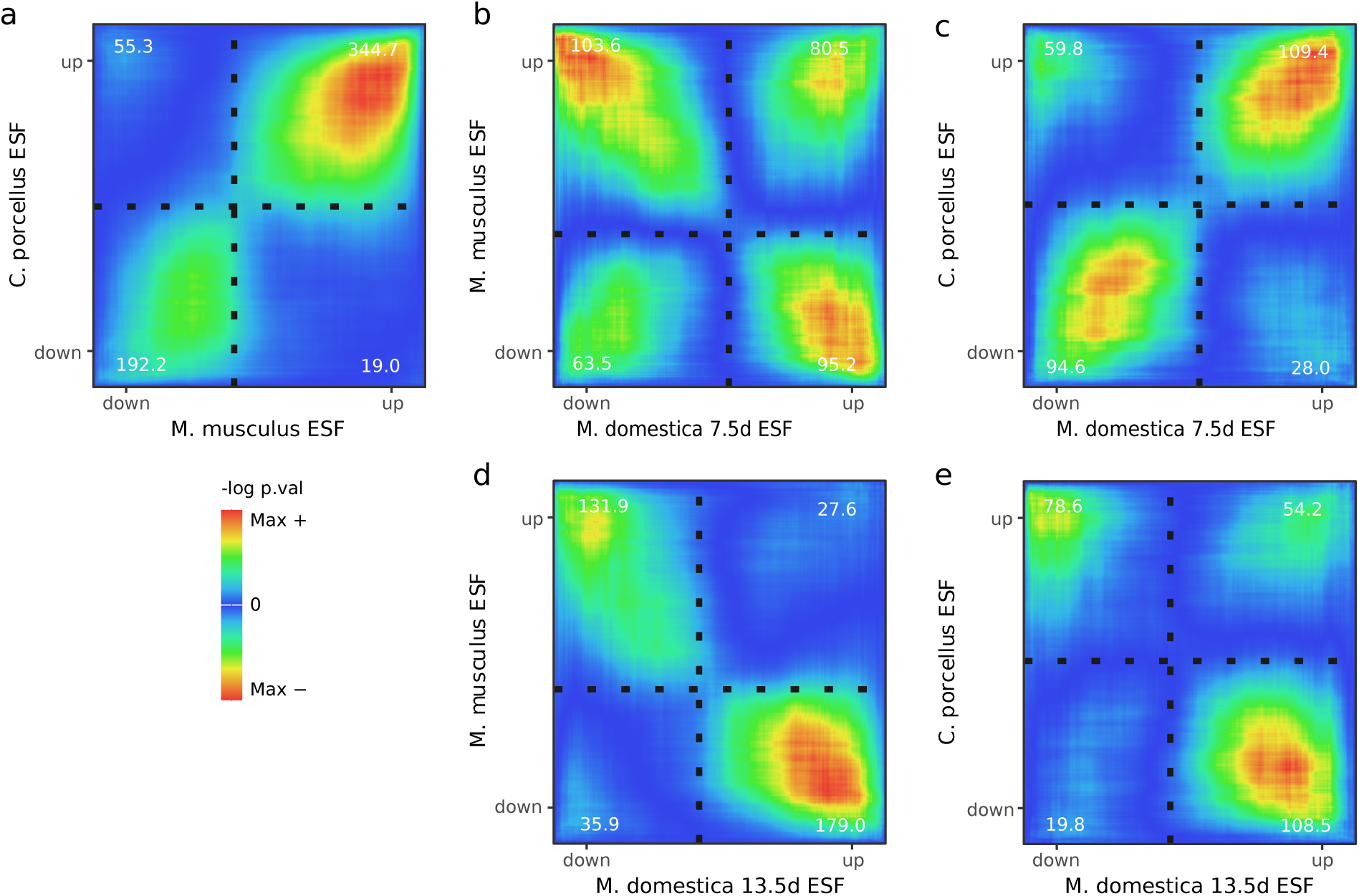
Concordance across species in gene expression changes during the transition from non-pregnant to peri-implantation. Cross-species rank-rank hypergeometric overlap (RRHO) heat maps compare differential gene expression test results between two paired comparisons. Horizontal and vertical axes represent paired genes, positioned in rank order of MAST differential expression test statistic of the same gene in both comparisons, assigned a positive or negative sign according to whether gene expression increases or decreases between the two stages compared. Pixels are colored according to the statistical significance of the overlap (-log of RedRibbon hypergeometric overlap test p-value) between the two expression changes under comparison, and quadrants are annotated with the highest value in the quadrant. Signal in the lower-left and the upper-right quadrants indicates concordant changes (in the same direction in both species), whereas the upper-left and the lower-right quadrants indicate discordant changes. **a-c**) Pairwise comparison between endometrial stromal fibroblasts (ESF) gene expression changes from the non-pregnant to peri-implantation stage of *M. musculus* and *C. porcellus* (**a**) *M. domestica* and *M. musculus* (**b**) and *M. domestica* and *C. porcellus* (**c**). **d-e**) Pairwise comparison between ESF gene expression changes from the non-pregnant to placentation (13.5 dpc) stage of *M. domestica* and the ESF gene expression changes from the non-pregnant to peri-implantation stages of *M. musculus* (**d**) and *C. porcellus* (**e**).

### Evolutionary changes in embryonic gene expression are insufficient to explain the origin of implantation

An alternative, embryo-centric model is that embryo implantation could conceivably have evolved via the early embryo and trophoblast gaining novel invasive or uterine-remodeling functions. To investigate this possibility, we analyzed single-cell transcriptomes from blastocysts stage-matched to our three species of mouse (E4.5) (52), guinea pig (E5.5) (53), and opossum (E7.5) (13), as well as the implantation-stage human (E7) (54). We focused our analysis specifically on functional genes curated from the literature with prior expectation to be involved in early blastocyst implantation and hatching (**Table S5**), and on cells from the trophectoderm, as the population that makes first contact with the uterine lining. Functional candidate genes included serine proteases, based on their roles in hatching from embryo coats and digestion of matrix to embed into the uterus (14, 16, 55, 56), enzymes catalyzing synthesis of steroid hormones such as estradiol and progesterone that affect uterine remodeling, and growth factors known to exert paracrine effects on the endometrium (57, 58).

Trophectoderm protease expression was found to be highly conserved across marsupial, rodent, and primate species, particularly *ADAM* family genes and cathepsins **(Figure 5a)**. Enzymes involved in steroid biosynthesis in the blastocyst showed conservation as well, including dehydrogenase enzymes involved in estradiol synthesis from cholesterol, such as *HSD17B7*, *HSD17B11,* and *HSD17B12* **(Figure 5a)** (59, 60). Trophectoderm production of growth factors such as *WNT3A*, *WNT6*, *BMP2,* and *BMP4*, and was also largely consistent across all four species. Some growth factors were only detected in eutherian embryos, including *LIF* (61). The embryos of therian mammals utilize a common molecular toolkit involved in apposition and hatching, but these do not partition by implantation mode and are therefore insufficient to explain the origin of implantation.

**Figure 5.**
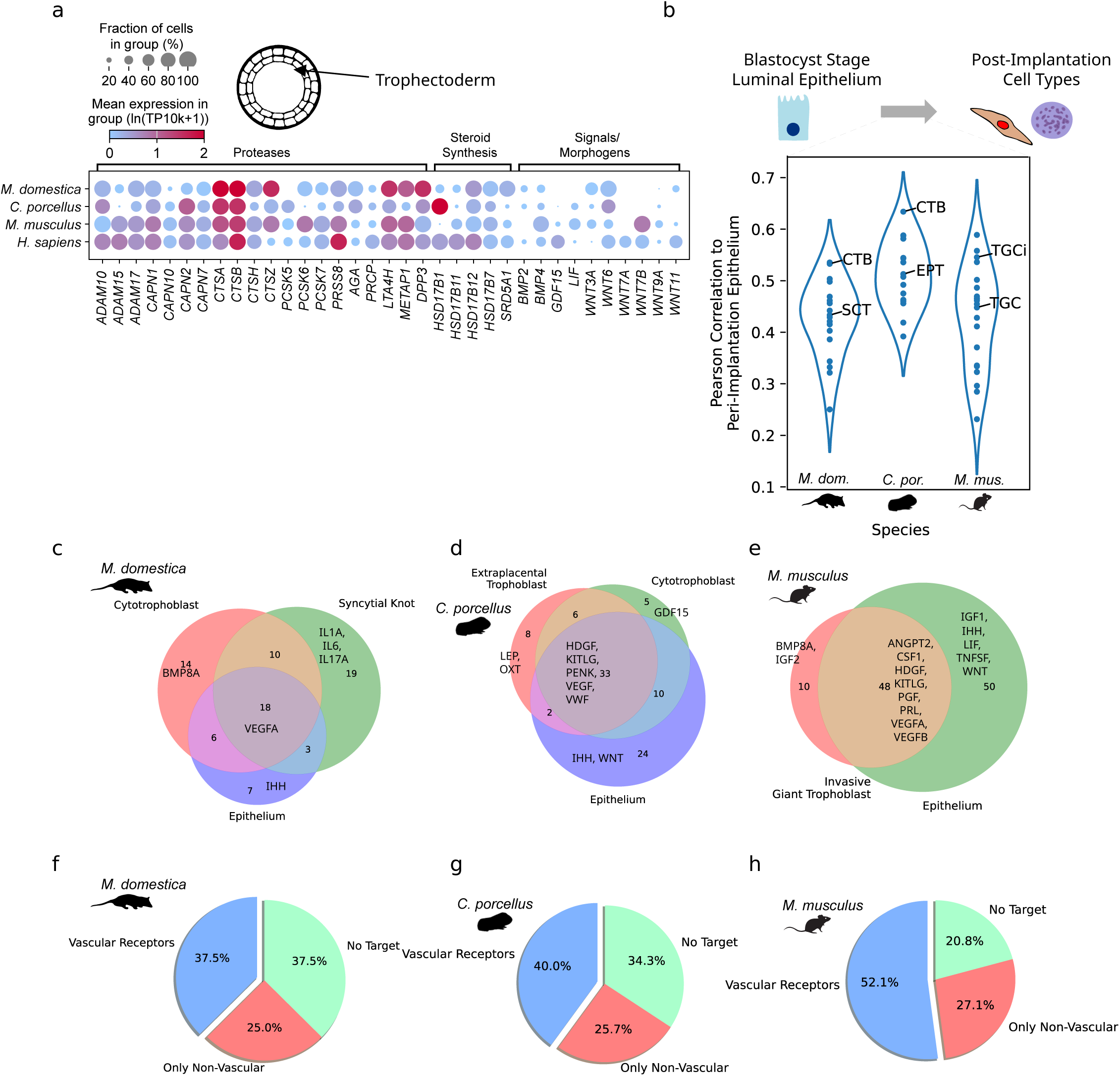
Conservation in trophectoderm and trophoblast functional gene expression. **a**) Expression of proteases, steroidogenic enzymes, and implantation-associated growth factors in trophectoderm cells from the opossum, guinea pig, mouse, and human implantation-stage blastocyst. **b**) Pearson correlations between signaling ligand gene expression in peri-implantation luminal epithelial cells with ligand gene expression of all cell types from the post-implantation stage of the same species. **c-e**) Venn diagrams of expressed ligand genes between peri-implantation epithelium and mid-gestation trophoblast. **f-h**) Pie charts showing the percentages of epithelial-trophoblast substituted ligands from (**b**) with corresponding receptors expressed only in vascular cell types (“Vascular Receptors”), only in non-vascular cell types (“Only Non-Vascular”), or with no corresponding receptor detected in the scRNA-seq data available (“No Target”). CTB: cytotrophoblast; SCT: syncytiotrophoblast; EPT: extraplacental trophoblast; TGCi: invasive trophoblast giant cell; TGC: trophoblast giant cell.

### Trophoblast substitutes for lost luminal epithelial signaling after invasion

One of the defining attributes of eutherian invasive implantation is the loss of the uterine luminal epithelium: in rodents, luminal epithelial cells undergo apoptosis and are phagocytosed by placenta cells soon after implantation occurs (62). How does the tissue remain stable after destroying the same cell type that earlier regulates stromal development and receptivity? Given the demonstrated importance of epithelial-stromal signaling to early implantation and its marsupial counterpart, we hypothesized that the loss of the luminal epithelium was compensated for by other cell types during the evolution of implantation. We re-analyzed a recent set post-implantation fetal-maternal interface transcriptomes from the mouse, guinea pig, and opossum (21) to test whether the paracrine signals once performed by the luminal epithelium are lost or maintained by other cell types after implantation.

We calculated the Pearson correlation between ligand-only transcriptomes of peri-implantation luminal epithelial cells and each annotated cell type in the mid-gestation (placentation) stage fetal-maternal interface (**Figure 5b**). Trophoblast cell types - the terminally-differentiated descendants of the blastocyst trophectoderm that make up the placenta - were among the most similar in ligand gene expression to the peri-implantation luminal epithelium in all species, including the non-invasive opossum. In the mouse, the top trophoblast cells corresponded to invasive (TGCi) and non-invasive (TGC) trophoblast giant cells; in both the guinea pig and opossum, the most epithelial-like cell type was fetal cytotrophoblast (CTB). Using an on/off threshold of 10% of cells in a cluster to binarize gene expression, we calculated the set intersections between ligand “secretomes” of these epithelial-trophoblast pairs (**Figure 5c-e**). Consistently overlapping signals between epithelium and trophoblast included cell type-specific growth factors (e.g. *VEGF*, *HDGF*). A substantial percentage of substituted ligands had receptors in the vasculature (52.1% in mouse, 40.0% in guinea pig, and 37.5% in opossum), while only 25-28% of substituted ligands targeted exclusively non-vascular cell types **(Figure 5f-h)**. Despite the fact that eutherian invasive implantation uniquely requires the destruction of the luminal epithelium, this pattern suggests that the trophoblast that takes its place is able to substitute the majority of the lost luminal epithelial cells’ signaling output, especially those targeted to the vasculature. Contrary to the prediction of compensation mechanisms being derived, this signaling similarity was detected even in our marsupial outgroup, and therefore was likely an ancestral feature of therian trophoblast that enabled, rather than responded to, the evolution of implantation.

## Discussion

To investigate how embryo implantation evolved, we compared two eutherians with invasive implantation, the mouse and guinea pig, with the opossum *Monodelphis domestica*, a marsupial that lacks implantation. We asked whether the evolutionary innovations enabling implantation occurred primarily in the embryo (the trophectoderm and trophoblast), the maternal tissue, or both. Our results support a model in which implantation evolved by modifying an ancestral progesterone-responsive endometrial remodeling program and coupling it to eutherian-specific developmental processes, including decidualization and epithelial erosion.

The marsupial and eutherian endometrium contain a similar repertoire of constituent cell types, but the hormone-driven remodeling of early pregnancy redirected them toward different tissue compositions. In the opossum, we observed a marked expansion of glandular epithelial cells at the shelled blastocyst stage that coincided with a nearly 300-fold upregulation of *IGF1* and estrogen-responsive glandular gene expression signature, a known regulator of *IGF1* (30). Indeed, comparative transcriptomics has suggested that suppression of estrogen signaling during the window of uterine receptivity is a derived trait of eutherians lacking in marsupials (63). Wnt signaling, important for glandular crypt formation, was upregulated in rodents but downregulated in the opossum. Functionally, these differences in cell type abundance and diversity likely correspond to the functional specifications of the two lineages: in eutherians the stroma develops into the decidua, with its invasion-regulating and immunoregulatory functions, whereas histotrophic nutrition in marsupials relies heavily upon glands. Glandular crypt invagination driven by Wnt signaling is necessary in rodents to form the chambers into which blastocysts implant (64), yet the histotroph they provide is less consequential than in marsupial gestation. Thus, the same epithelial and stromal cell types show dynamics that reflect the two lineages’ distinct developmental modes.

A central question in comparing marsupial and eutherian pregnancy is which marsupial stage, if any, corresponds to the eutherian window of receptivity. Our finding of a common cell-signaling cascade in early pregnancy of marsupials and eutherians suggests a shared foundation upon which both the eutherian implantation reaction and the marsupial attachment response were built (**Figure 6**). Single-cell transcriptomic analysis revealed greater similarity of the opossum uterus at the shelled-blastocyst stage (7.5 dpc) to the rodent peri-implantation uterus (mouse 4.5 dpc, guinea pig 6.5 dpc) than of the opossum’s placentation stage (13.5 dpc). This early stage resembles the eutherian window of receptivity because of the developmental stage of the embryo, a homologous maternal endocrine state as a result of ovarian progesterone (65), and progesterone/SOX17-IHH-BMP2 signaling activity. At this stage, we observe gene expression dynamics that resemble the progesterone-driven stages of rodent implantation involved in relaying epithelial IHH into decidualization. These observations suggest that the progesterone-driven first step of the implantation cascade was already functional in the uterus of the last common ancestor of marsupials and eutherians (**Figure 6c**), as was the second stage of stromal BMP2 expression. Yet, unlike in eutherians, its signaling output appears to be inhibited. We find a role for epithelial BMP2 inhibition via glandular *GREM2* - increasing from 2 to more than 2000 TPM - in the 7.5 dpc opossum endometrium. GREM2 is a secreted antagonist that binds BMP2 with high affinity (66) and shows high selectivity for BMP2 and BMP4 (49). Experimental disruption of *Bmp2* in mice leads to a phenotype where embryos still superficially attach - like in marsupials - but decidualization and invasion fail (67). The parallels of this to the marsupial condition warrant further investigation. In human and rodent endometrial stromal cells, decidualization in response to progestin and cyclic AMP occurs through a conserved BMP2-WNT4/β-catenin pathway (48, 68), and disruption of BMP2 signaling has been shown to lead to female-factor infertility due to decidualization failure in human endometriosis (69). Reduced stromal *BMP4* expression has been observed in both mouse embryo resorption and human recurrent implantation failure (70). In contrast, treatment of opossum endometrial stromal fibroblasts with eutherian deciduogenic stimuli upstream of BMP2 (cyclic AMP or PGE_2_ and progestin) induces oxidative stress and apoptosis (51). We therefore hypothesize that inhibition of the later steps in this pathway via *GREM2* may perform a cytoprotective function in the opossum that became unnecessary once the decidual cell type evolved in Eutheria with its extensive robustness to oxidative stress. Stromal competence to conjoin the output from the ancestral epithelial-stromal signaling axis with decidual differentiation was evidently a eutherian innovation.

**Figure 6.**
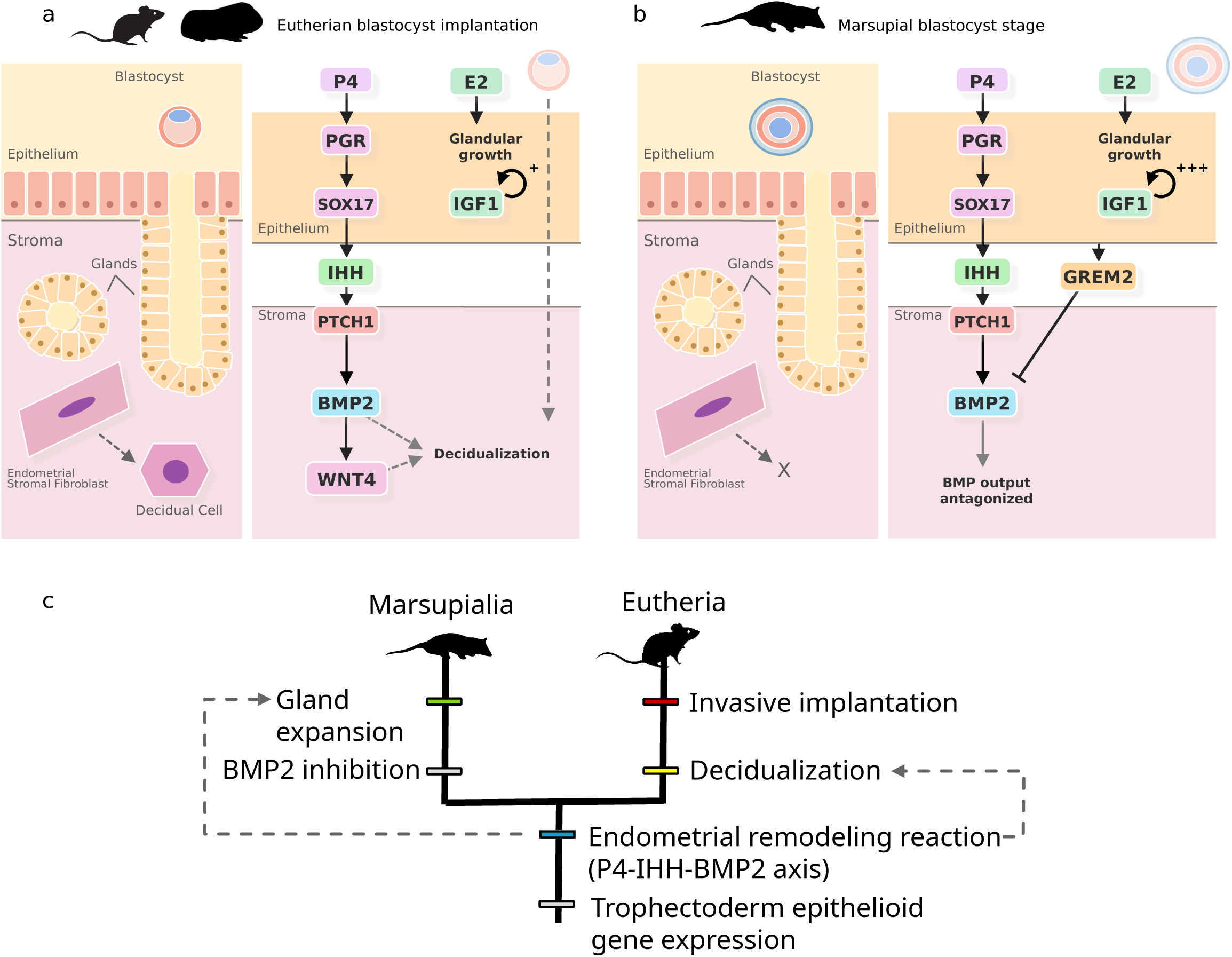
Model for the evolution of the implantation reaction. Schematized depiction of the corresponding uterine stage to the blastocyst-stage embryo in eutherian (rodent) peri-implantation (**a**) and the marsupial (opossum) shelled blastocyst stage (**b**). The right side of each panel shows the epithelial-stromal IHH signaling axis as it exists in each lineage. (**c**) Cladogram depicting the inferred order of evolutionary steps. Trophoblast epithelioid gene expression originated before the ancestral therian endometrial remodeling reaction (P4-IHH-BMP2 axis), which in turn was the basis for the eutherian implantation reaction and its counterpart in marsupial pregnancy. P4: progesterone.

*LIF* signaling from the glands was not detected in the opossum or guinea pig blastocyst-stage uterus. Studies in rodents suggest that the role of *LIF* in the window of implantation remodeling cascade is supplementary to progesterone, rather than essential (71). We therefore conclude that the core of the ancestral endometrial remodeling response - conserved between eutherians and marsupials - is elicited by progesterone, with additional modulatory factors having arisen secondarily.

We did not find strong support for an embryo-centric evolutionary scenario for the origin of implantation. Recent single-cell atlases of early embryonic development have failed to reveal consistent transcriptomic differences associated with implantation type (72). We analyzed time-matched early-embryo cell atlases for our three species and the human to test for gene-expression differences across the marsupial-eutherian divide in pathways functionally relevant to implantation. Rather than a marsupial-eutherian divergence, we found a high degree of conservation in pathways including matrix and shell coat-degrading proteases, steroidogenic enzymes, and growth factors whose receptors are expressed in the endometrium. This conservation suggests that the signaling potential of the blastocyst did not undergo a dramatic change in these implantation-related gene expression pathways unique with eutherians. While our analysis was not designed to exhaustively identify embryo-specific adaptations, these findings are consistent with the maternal-first hypothesis for the origin of embryo implantation (12), in which changes in maternal tissue architecture and responsiveness first established the developmental context to support invasive placentation.

Our comparison of the secreted signaling repertoire of the uterine epithelium at peri-implantation and of trophoblast at mid-gestation revealed continuity of ligand-receptor interactions across pre-implantation to post-implantation stages of pregnancy. Despite the loss of epithelial cells in eutherians, the majority of the lost luminal epithelial signals are substituted by the post-implantation trophoblast, and quantitative comparison of cell type secretomes revealed that trophoblast consistently bore the greatest similarity to the lost luminal epithelium . We term this phenomenon “cell type substitution”: the replacement of one member in an established cell-signaling network by another cell type that preserves key signaling relationships. Redundancy between cytotrophoblast-secreted ligands and those of the uterine luminal epithelium was also apparent in the opossum. The capacity of the trophoblast to substitute for luminal epithelial signaling likely did not evolve to facilitate implantation, but rather stems from the conserved epithelial-like nature of trophoblast cell types, which derive from ectodermal progenitors. We propose that eutherians co-opted this property of the ancestral therian trophectoderm during the evolution of invasive placentation.

Invasive implantation is the defining innovation of eutherian pregnancy. Previous work has established the decidual stromal cell type as a key innovation behind eutherian pregnancy, produced through gene-regulatory rewiring of ancestral endometrial fibroblasts (73) and the gain of cell type-specific functions supporting invasive placentation (74). Our results help to place that innovation in a broader developmental context. The opossum does not possess a decidual stromal cell type, but shares a surprising degree of the upstream epithelial-stromal signaling axis that, in eutherians such as rodents, feeds into embryo-induced decidualization. The evolution of implantation required rewiring of the maternal signaling state so that the ancestral remodeling program could transition into decidualization, stable luminal epithelial loss, and invasion. Future work will be needed to reconstruct, experimentally and through profiling of species at more pivotal phylogenetic nodes, how this conserved uterine signaling axis became coupled to a new stromal cell type and thereby enabled an intimate and prolonged gestation.

## Materials and Methods

### Sample Collection and Design

In all three species, we targeted the stage when the peri-implantation embryo is a blastocyst, and maternal serum progesterone is elevated: 4.5 dpc for the mouse (75), 6.5 dpc for the guinea pig (76), and two stages of the opossum: 7.5 dpc, when the fetus is still a blastocyst (comparable to the peri-implantation blastocyst in rodents) surrounded by the soft shell coat (77, 78), and day 13.5 of gestation, when the shell coat surrounding the fetus is lost and placental attachment has begun.

*C. porcellus* (Charles River) were maintained at the University of Vienna according to Institutional Animal Care protocols. The estrous cycle was monitored by examination of vaginal membrane opening following (79). Females were mated while in estrus at 3-4 months of age, and video recording was used to detect copulation during the night. Two individuals were used for sampling the implantation stage, 6.5 dpc and one was used as non-pregnant control (at 6 days post estrus, i.e. early diestrus).

*M.musculus* (C57BL/6J) were maintained at the University of Vienna according to Institutional Animal Care protocols in a separate facility. The estrus cycle was monitored by vaginal swabbing following (80). Copulation was determined by the presence of a copulatory plug and considered as day 0.5 post-copulation. Two individuals were used for sampling the implantation stage at 4.5 dpc and two were used as non-pregnant controls (at 2 days post estrus, i.e. early diestrus, as staged by vaginal cytology).

*M. domestica* were raised in a breeding colony at Yale University according to ethical protocols approved by the Yale University Institutional Animal Care and Use Committee (#2020-11313). Two individuals were used for sampling each stage: the non-pregnant/anestrus endometrium (n=2), 7.5 dpc (n=2) and 13.5 dpc (n=2, previously reported in (21)). Video recording was used to assess the precise time of copulation. If multiple copulations were observed, the first was used to establish 0 dpc.

### Single-Cell Dissociation

Whole uterine horns were dissected into phosphate-buffered saline (PBS). Portions of ∼ 0.2g per individual were minced into ∼1 mm^3^ cubes and transferred into a digestive solution containing 0.2 mg/mL Liberase TL (05401020001, Sigma) in 1800 μL PBS. The tissue was then incubated at 37°C for 15 minutes and then passed 10 times through a 16-gauge needle attached to a 3-mL syringe. This process was repeated another two more times, the last time with a 18-gauge needle for complete dissociation. 2 mL of charcoal-stripped fetal bovine serum (100-199, Gemini) were added to stop digestion by inverting the tube several times. After that, the cell suspension was passed through a 70-μm cell strainer then a 40-μm cell strainer to get rid of any remaining chunks of tissue. The filtered cell suspension collected was centrifuged at 300 g for 5 minutes. The resulting pellet was resuspended in 1x ACK red blood cell lysis buffer (A1049201, Thermo-Fisher), incubated at room temperature for 5 minutes, and centrifuged again. The final pellet was resuspended in PBS containing 0.04% bovine serum albumen (A9647, Sigma) and Accumax (07921, Stem Cell Technologies). The resulting single cell suspension was assessed with a Cellometer (mouse, guinea pig) or hemacytometer (opossum) to assess cell concentration and viability with trypan blue stain. Only cell suspensions with viability higher than 80-85% (rodent) or 70% (opossum) were used.

### Library Preparation and Sequencing

Cells were captured using the 10x Genomics Chromium platform (3’ chemistry, version 3). Mouse and guinea pig libraries were prepared at the University of Vienna, and opossum libraries were prepared at the Yale Center for Genomic Analysis following identical protocols (CG000315) from 10x Genomics. Libraries were sequenced using an Illumina NovaSeq 6000 by the Yale Center for Genomic Analysis (*M. domestica*) at a read depth exceeding 20,000 reads/cell and at the Next Generation Sequencing Facility of the Vienna Biocenter (*C. porcellus* and *M. musculus*) at a read depth of around 400M reads/sample (Illumina NovaSeq S4 PE150 XP for *C. porcellus* and Illumina NovaSeq SP Assymetric 10X for *M. musculus*).

### Single-Cell Data Analysis

Sequencing reads were aligned to reference genomes using the 10x Genomics CellRanger software (≥v7.0.0). *Monodelphis domestica*, *Cavia porcellus*, and *Mus musculus*, were mapped to their respective Ensembl genome annotations (ASM229v1 v104, cavPor3.0 v104, and GRCm39 v104, respectively). Feature count matrices were processed using CellBender remove-background (v0.3.2) (81) to remove background contamination before cell type annotation.

Species-specific single-cell datasets were analyzed and annotated separately following scanpy standard functions and criteria to reveal species-specific cell types. Cells with fewer than 700 unique features or greater than 25% of transcripts of mitochondrial origin were filtered, as well as cells predicted to be doublets by doubletdetection (v4.2) (82). In total, 24,752 cells from the mouse, 29,750 from the guinea pig, and 43,561 from the opossum were retained. Library size normalization, log1p normalization, feature selection, dimensionality reduction, clustering and marker gene identification were performed using scanpy ≥ v1.9.1 (83). Replicates belonging to the same species were corrected for batch effect using harmony (harmonypy, v0.0.9) (84), which adjusts principal components to aid in cluster delimitation but does not alter expression values.

Time-matched peri-implantation blastocyst scRNA-seq data were obtained from public repositories. These included opossum E7.5 bilaminar blastocyst (13), mouse E4.5 blastocysts (52), guinea pig E5.5 blastocysts (53), and human day 7 blastocysts (54). Blastocyst cell types were annotated to identify trophectoderm using the original authors’ annotations in the case of opossum and human. For mouse and guinea pig, where the authors did not provide original cell annotations, trophectoderm was identified as a *CDX2* and *GATA3*-expressing cell population following dimensional reduction and leiden clustering. Post-implantation scRNA-seq data of the mouse and guinea-pig and 13.5 dpc opossum were obtained from (21) with original annotations preserved, and processed with CellBender remove-background as above for methods consistency.

Non-pregnant and early pregnancy datasets from all species were integrated into a shared UMAP manifold using SAMap (v.1.3.4) (23). Pairwise mapping scores between putative cell type clusters across species were calculated using the get_mapping_scores() function on the pooled transcriptome of all cells in each cluster. Confidence intervals for SAMap mapping scores were generated using the bootstrapping method described in (21), in which mapping scores were re-calculated on 200 random 30% subsamples of the full SAMap graph, and the bootstrapped mean score of all cell type pairs over the 200 iterations was used for plotting.

### Differential Gene Expression Analysis

To identify differentially expressed ligand and receptor signaling from epithelial to stromal cells at implantation, differential gene expression analysis was performed using MAST (v3.20) (85). Significantly changed genes were classified as those with a Benjamini-Hochberg adjusted MAST test p-values of less than 0.05. To identify differentially expressed ligands and receptors, the list of genes subjected to differential expression testing was subset to only genes encoding ligands, receptors, or the subunits of either in our ligand-receptor ground truth database used for communication inference. Gene set enrichment on differentially expressed genes was conducted using GSEApy (v1.1.3) (86) against the gene sets of single-molecule perturbations on cultured cell lines available in the NCBI gene expression omnibus (Enrichr “Gene_Perturbations_from_GEO_up”).

To compare differential expression responses in select cell types between non-pregnant and blastocyst-stage uteri across species, we performed rank-rank hypergeometric overlap analysis using the RedRibbon R package (v1.3-1) (50). For each species pair and cell type tested, we used the single-cell MAST differential expression output. To ensure comparable gene sets, genes were subset to those annotated by Ensembl as one-to-one orthologs. Genes were ranked by the MAST regression coefficient (“coef”) to preserve directionality of differential expression and supplied to the RedRibbon() function using the “hyper-two-tailed” (two-tailed hypergeometric enrichment) mode to assess overlap. Results were passed to the quadrants() function to extract overlapping gene sets using the “ea” (evolutionary algorithm) mode and permutation-based hypergeometric test p-value adjustment.

### Cell-Cell Communication Analysis

We inferred cell communication events between cell type transcriptomes using a method adapted from (21). First, a ground truth ligand-receptor database was adapted from the manually-curated fork of CellPhoneDB v5.0.0 (87) used in (21). To remove ambiguity from multiple-subunit receptor complexes and allow 1:1 differential expression testing, the ground truth database was modified to include only the subunit that actively binds the ligand (excluding co-receptors), or if multiple, to include each as a separate ligand-receptor pair. Genes were mapped to their one-to-one human ortholog according to the Ensembl database orthology, or else excluded. Cell interactions were inferred using expression thresholding with a cutoff of 0.2 (20% of cells in the cluster) for both ligand and receptor using chinpy (v0.0.55; https://gitlab.com/dnjst/chinpy). Statistical testing for significantly cell type-enriched interactions was conducted using LIANA+ (v1.2.0) (37) with the same parameters, and the outputs were merged for plotting and reporting.

For circular plotting of inferred ligand-receptor interactions, depicted interactions were subset to those annotated as “Secreted Signaling” in our database (paracrine and endocrine secreted ligands) and “Small Molecule-Mediated” (enzymes producing steroid hormones and other small molecules), but excluding extracellular matrix-mediated ligand-receptor interactions, those requiring direct cell-cell contact, and those with questionable ligand status; an expanded list of inferred interactions and test statistics behind the circular plots is reported in **Table S4**.

For analysis of epithelial cell substitution, Pearson correlations were calculated between the peri-implantation uterine epithelium signaling repertoire (this study) and the mid-gestation secreted signal repertoires of all cell types from the corresponding species dataset (21) using the corr() function of the pandas (v2.2.2) package. Input data were boolean matrices of all ligands classified as Secreted Signaling (a total of 302), with a threshold proportion of cells in the cluster to be considered “on” of 0.10. Venn diagrams were plotted using the matplotlib_venn (v1.1.1) package.

## Acknowledgments

The authors thank the Yale Center for Genomic Analysis (NIH 1S10OD030363-01A1) for library preparation and use of their high-performance computing cluster.

## Data, Materials, and Software Availability

Sequencing data for peri-implantation as well as diestrus/non-pregnant uteri of mouse, guinea pig, and opossum are deposited in the NCBI Gene Expression Omnibus at GSE292958. Mid-gestation stage utero-placental interfaces of the same species were re-analyzed from data deposited at GSE274701. Opossum day 7.5 blastocyst data are available as EMBL ArrayExpress E-MTAB-7515, guinea pig blastocyst transcriptomes were accessed from China National Center for Bioinformation PRJCA028188 , mouse day 4.5 blastocyst transcriptomes were accessed from NCBI GEO GSE63266; and human day 7 blastocyst at EMBL ArrayExpress E-MTAB-3929. No software was developed specifically for this study. Analysis notebooks are archived at https://gitlab.com/dnjst/implantation26.

## Author Contributions

Conceptualization: MP, GPW

Funding acquisition: MP, GPW, DJS

Experimentation: SB, DJS, CMG, JDM, AGC

Animal colony maintenance: JDM, DJS, SB, GD

Data analysis: DJS, SB

Supervision: MP, GPW

Writing - original draft: SB, DJS

Writing - review & editing: SB, DJS, GPW, MP, CMG

## Preprint Servers

This manuscript has been deposited on BioRxiv under a CC-BY-ND 4.0 International license at https://doi.org/10.1101/2024.11.04.621939.

---

**Figure S1.**
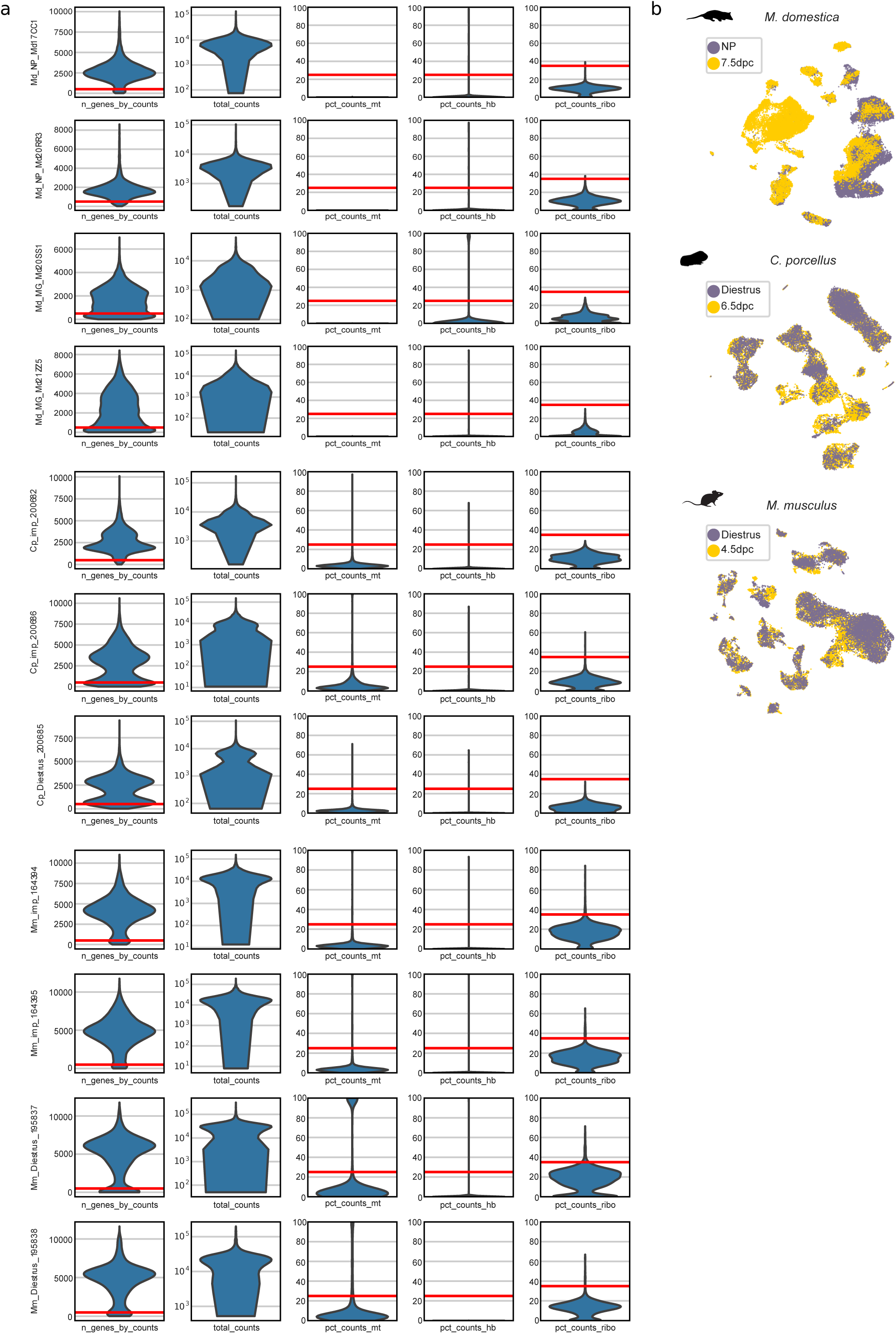
Quality control statistics for single-cell data. **a**) Violin plots showing quality control distributions and cutoffs (red lines) for numbers of genes per cell (n_genes_by_counts), total transcripts per cell (total_counts), % mitochondrial counts (pct_counts_mt), % hemoglobin counts (pct_counts_hb), and % ribosomal gene counts (pct_counts_ribo) per library. Md: *M. domestica*; Cp: *C. porcellus*; Mm: *M. musculus*. **b**) Within-species UMAP embeddings after batch integration (harmony), colored by stage.

**Figure S2.**
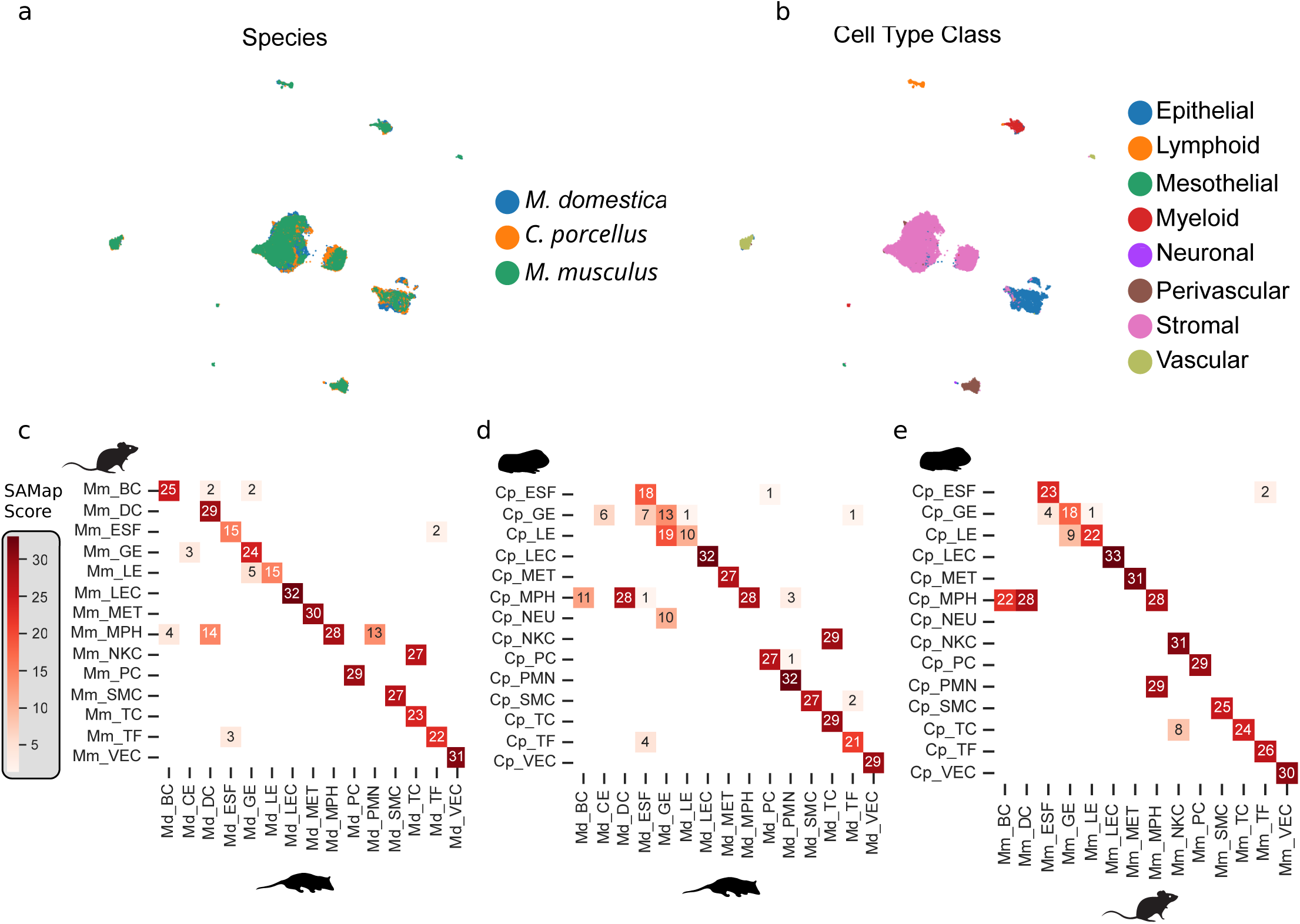
SAMap integration. **a**) Integrated embedding colored by species. **b**) Integrated embedding colored by cell type class. **c-e**) Heatmaps showing bootstrapped pairwise SAMap transcriptomic similarity scores (multiplied by 100) between pairs of annotated clusters in each species. Md: *Monodelphis domestica*; Mm: *Mus musculus*; Cp: *Cavia porcellus*.

**Figure S3.**
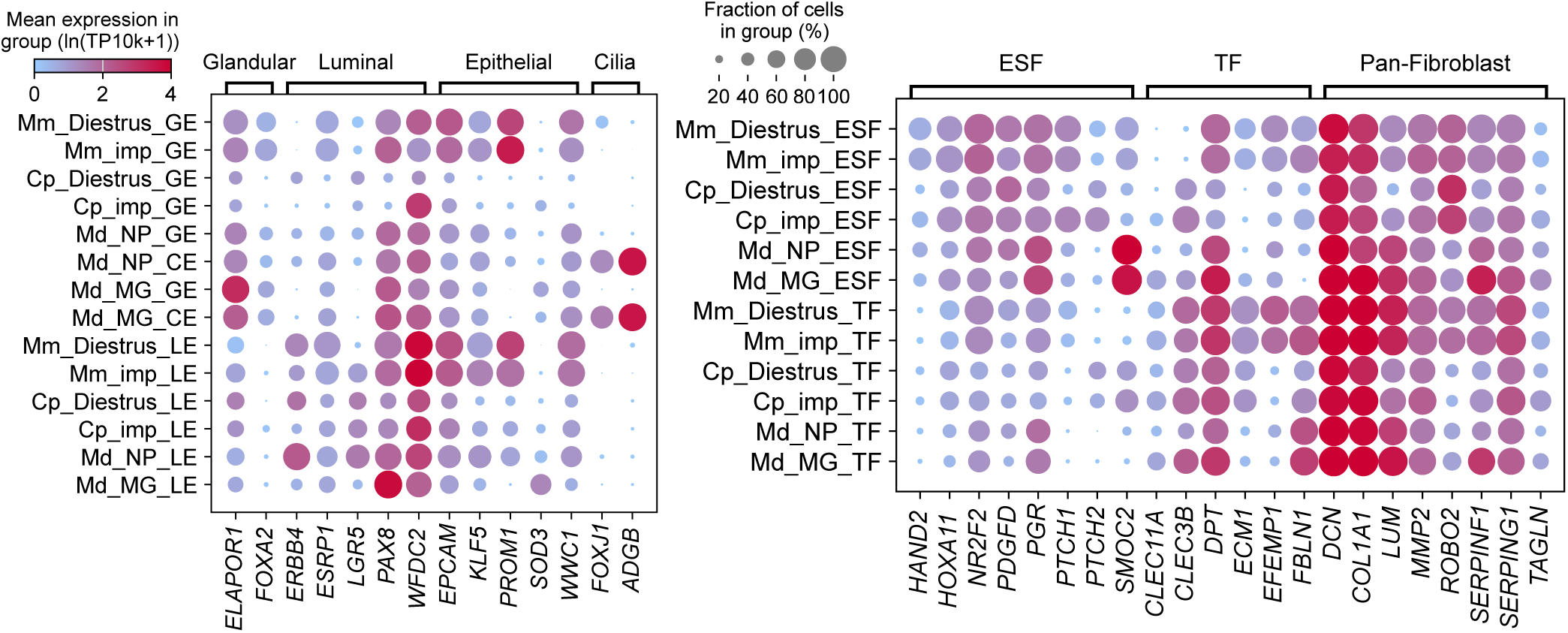
Additional marker gene expression dotplots. Selected mouse (Mm), guinea pig (Cp) and opossum (Md) epithelial and stromal cell type maker genes at implantation (imp), diestrus and mid-gestation (MG), and non-pregnant (NP) stages. GE: glandular epithelium; LE: luminal epithelium; CE: ciliated epithelium; ESF: endometrial stromal fibroblast; TF: myometrial tissue fibroblast.

**Figure S4.**
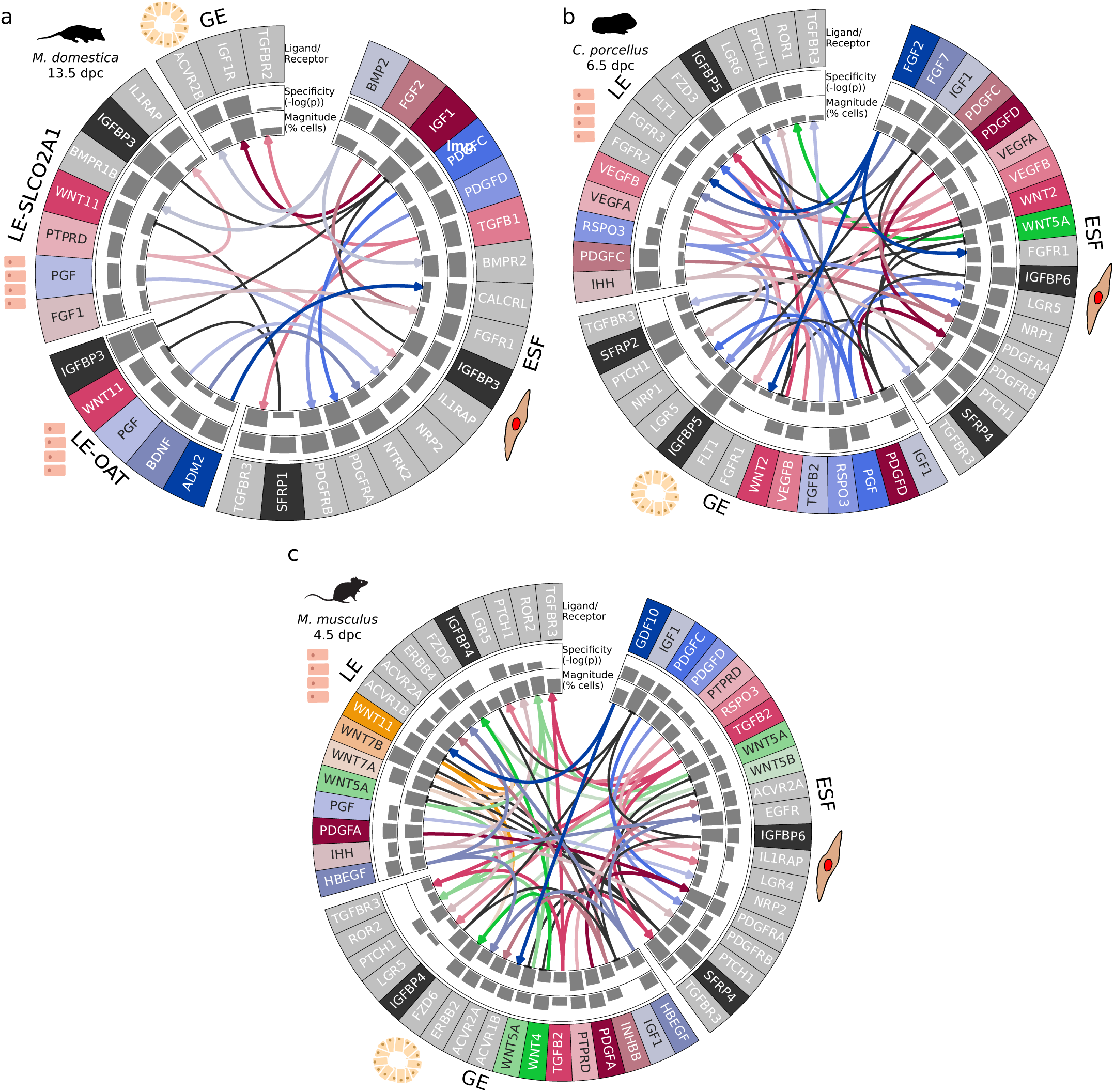
Additional ligand-receptor interaction networks between epithelial and stromal cell types. Arrows show directions of inferred ligand-receptor signaling. Outer barplots show LIANA+ rank-aggregate specificity score, while inner barplots show the percent of cells in the cluster expressing the ligand or receptor. Ligands are shown in color, receptors in grey, and inhibitors in black. ESF: endometrial stromal fibroblast; GE: glandular epithelial cell; LE: luminal epithelial cell; LE-SLCO2A1: SLCO2A1+ luminal epithelial cell; LE-OAT: OAT+ luminal epithelial cell.

